# Cell cycle progression requires repression by Groucho during S-phase and its relief at G2-phase

**DOI:** 10.1101/2022.02.22.481413

**Authors:** Shaked Bar-Cohen, Ze’ev Paroush

## Abstract

The cell cycle depends on a sequence of steps that are triggered and terminated via the synthesis and degradation of phase-specific transcripts and proteins. While much is known about how stage-specific transcription is activated, less is understood about how inappropriate gene expression is suppressed. In this paper we demonstrate that Groucho, the *Drosophila* orthologue of TLE1 and other related human transcriptional corepressors, regulates cell cycle progression *in vivo*. We show that although Groucho is expressed throughout the cell cycle, its activity is selectively inactivated by phosphorylation, except during S-phase when it represses *e2f1* expression. Misregulated Groucho activity causes cell cycle arrest; in particular, both constitutive Groucho activity and failure to repress *e2f1* cause cell cycle arrest phenotypes. We also show that the Cdk1 kinase is responsible for stage-specific phosphorylation of Groucho *in vivo*. We propose that Groucho and its orthologues play key roles in the metazoan cell cycle that may explain the links between TLE corepressors and several types of human cancer.

## Introduction

The cell cycle is composed of a programmed sequence of events, including DNA synthesis, chromosome separation and cytokinesis. Progression through the cycle is highly regulated by protective checkpoints at which intrinsic or extrinsic conditions are monitored. The cycle arrests if suboptimal conditions are sensed, and recommences only once these are resolved through appropriate cellular response mechanisms (Clarke and Giménez-Abián, 2000; Kastan and Bartek, 2004; Musacchio and Salmon, 2007).

Cell cycle-related proteins are regulated at different levels. In the early *Drosophila* embryo, before the maternal-to-zygotic transition, rapid nuclear divisions are mainly controlled via translational and post-translational regulation of maternaly-deposited determinants (Becker et al., 2018; Groisman et al., 2002). As zygotic transcription is induced later in development, additional mechanisms are incorporated as examplified by the surge of *string* (*cdc25*) expression that is responsible for introducing the Gap 2 (G2)-phase in otherwise Synthesis (S)-to-Mitosis (M) cycling embryonic cells (Budirahardja and Gönczy, 2009; Edgar et al., 1994). Other genes that play a part in the cell cycle are also transcriptionally regulated, e.g., those involved in processes such as DNA replication and chromosome segregation, many of which are highly expressed in human tumors (Whitfield et al., 2002). Thus, transcriptional regulation in the contex of the cell cycle is an important path to explore.

E2F1, a key cell cycle transcription factor, is an activator of *cyclin E* (*cycE*) expression, as well as that of additional target genes whose protein products are required at the initiation of S-phase, and at the G2- and M-phases (Bennett et al., 1996; Dimova and Dyson, 2005; Dimova et al., 2003; Herr et al., 2012). The E2F1 protein is detectable during all stages of the cell cycle except for S-phase, when it is degraded (Davidson and Duronio, 2012; Reis and Edgar, 2004; Shibutani et al., 2008). Preventing E2F1 protein degradation during S-phase leads to acceleration of the cell cycle and/or to apoptosis, highlighting the requirement for its removal at this stage (Shibutani et al., 2008). The activator functions of E2F1 and other cell cycle factors have been studied in depth; less is known, however, about transcriptional repressors in this process.

We have previously shown that Transducin-like enhancer of split 1 (TLE1), a human orthologue of the *Drosophila* developmental corepressor Groucho (Gro), acts as an anti-proliferative factor (Zahavi et al., 2017), and TLE family members have been linked to human cancers (Dali et al., 2018; Kokabu et al., 2017). Furthermore, Gro and TLE1 are both phosphorylated in a cell cycle-regulated manner *in vitro* and in cultured cells (Nuthall et al., 2002), a modification previously shown to mitigate their corepressor activity downstream of receptor tyrosine kinase (RTK) pathways (Cinnamon et al., 2008; Hasson et al., 2005; Helman et al., 2011; Helman et al., 2012; Johnston et al., 2016; Nuthall et al., 2002; Paroush et al., 1997; Zahavi et al., 2017). Herein, we explore the possibility that Gro fulfils an *in vivo* regulatory function in the cell cycle.

We now demonstrate that Gro-mediated repression at S-phase, and its relief at G2-phase, are both critical for cell cycle progression. Specifically, we show that Gro is unphosphorylated and therefore active as a repressor only at S-phase, when it represses *e2f1* expression. *gro-*deficient cells display accelerated cell cycles and accumulate at Gap 1(G1)-phase, a phenotype resembling that caused by E2F1 overexpression. In addition, we find that overexpression of Gro leads to a G2-phase arrest, and identify phosphorylation by Cyclin-dependent kinase 1 (Cdk1) as a mechanism that attenuates Gro’s repressor activity at this stage *in vivo*, thus permitting entry into mitosis. Together, our results reveal an essential role for Gro in the cell cycle, showing that it switches between active and inactive states and restricts gene expression in a phase-specific manner. We propose that a similar level of regulation underlies the involvement of TLE corepressors in cancer.

## Results

### Groucho is selectively phosphorylated and inactive at mitosis and in the two Gap phases

Gro is uniformly expressed throughout *Drosophila* development (Delidakis et al., 1991). In any given nucleus, however, it is either fully phosphorylated or completely unmodified (Cinnamon et al., 2008; Helman et al., 2011; Johnston et al., 2016) (Suppl. Fig. S1; see Materials and Methods). In previous work we demonstrated that phosphorylation of Gro, particularly by extracellular signal-regulated kinase (Erk) in response to RTK signaling, downregulates its repressor function (Cinnamon et al., 2008; Hasson et al., 2005; Helman et al., 2011; Helman et al., 2012). Thus, Gro has a regulated ability to switch between two modes: an active (unphosphorylated) or an inactive (phosphorylated) corepressor.

Staining of cycling cells in wing and eye imaginal discs with antibodies specific for Gro and phosphorylated Gro (pGro; raised using a synthetic phosphopeptide containing one of two putative Erk consensus sites in Gro) revealed that they display minimal overlap in their respective signals; only 10% of cells in wing imaginal discs (n=1031) and 11% of cells in eye imaginal discs (n=492) show overlapping nuclear signals for both antibodies (Suppl. Fig. S1; see Materials and Methods) (Cinnamon et al., 2008; Delidakis et al., 1991; Hasson et al., 2005; Helman et al., 2011; Johnston et al., 2016). Using these two antisera, we find that phosphorylation of Gro fluctuates dynamically in a cell cycle phase-dependent manner, and that it is modified at all stages of the cell cycle except for S-phase.

Specifically, mitotic cells, distinguishable by anti-phospho-Histone 3 at Serine 10 (pH3) staining, are consistently positive for pGro, both in embryos (Fig. 1a-b’) as well as in wing imaginal discs (Fig. 1e-f’; quantified in Fig. 1i). In contrast, the pH3 signal complements that of the unphosphorylated, active form of Gro in both tissues (Fig. 1c-d’ and Fig. 1g-h’; quantified in Fig. 1i), indicating that Gro is phosphorylated, and therefore inactive as a repressor, at mitosis.

**Fig. 1.**
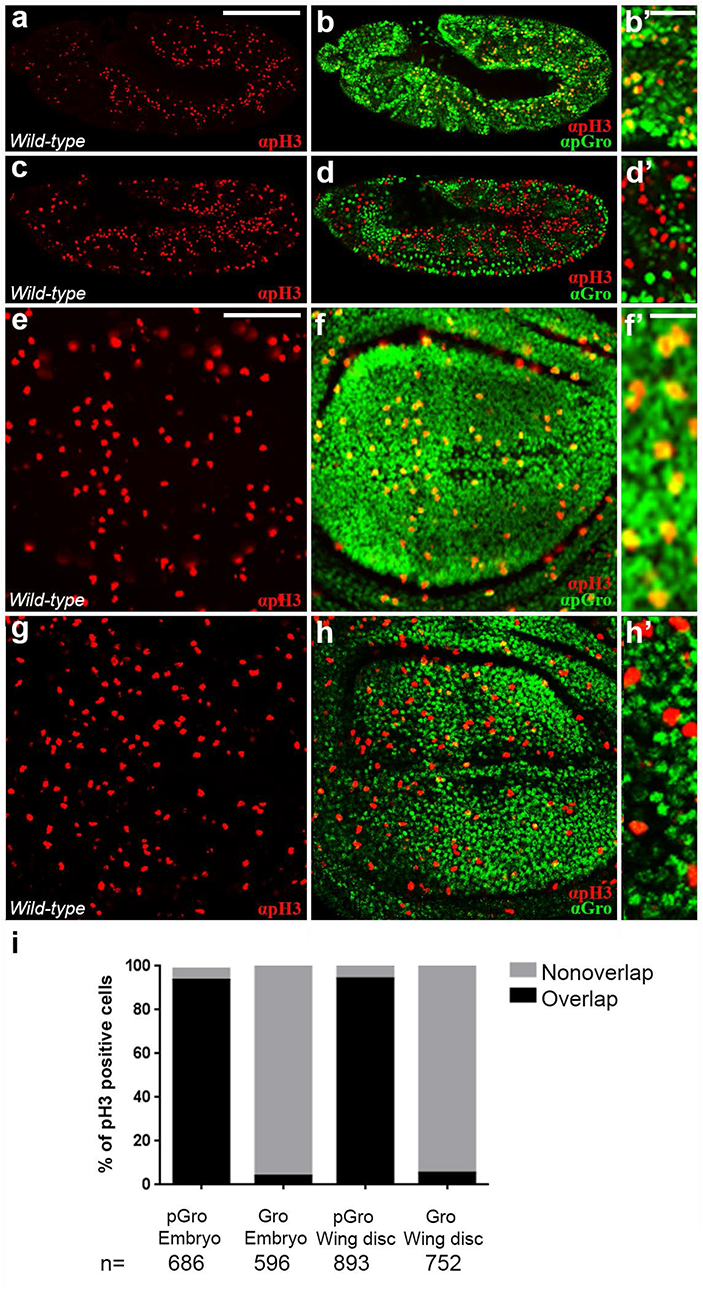
Groucho is phosphorylated in pH3-positive mitotic cells. (a-h’) Confocal images of stage 11 *wild-type* embryos (lateral views; a-d’) and wing imaginal discs from *wild-type* third instar wandering larvae (e-h’), co-stained for pH3 (red; a-h’) together with either pGro (green; b-b’ and f-f’) or Gro (green; d-d’ and h-h’). (b’, d’, f’, h’) Magnified views of (b, d, f and h), respectivley. In both tissues, pH3 staining overlaps extensively with that of pGro (b-b’ and f-f’) and complements that of Gro (d-d’ and h-h’). (i) Percentage of pH3-positive nuclei co-stained (black), or not (grey), with pGro or with Gro in embryos (two left columns) and in wing imaginal discs (two right columns). n = number of pH3-positive cells scored in each case. Scale bars = 100 µm (a-h) and 16.6 µm (b’, d’, f’, h’).

Not all pGro positive nuclei are mitotically active, however; we find that Gro is also phosphorylated at the G1- and G2-phases as determined using *Fly-FUCCI*, an *in vivo* fluorescent, ubiquitination-based indicator system (Fig. 2) (Zielke et al., 2014). In *Fly-FUCCI* wing imaginal discs, individual cells appear in different colours according to their cell cycle stage (Fig. 2a-b’, e-e’) (Zielke et al., 2014). Staining with anti-pGro and anti-Gro antibodies demonstrates that Gro is also phosphorylated at the G2/M- and G1-phases (Fig. 2c-d’, f-g’). We surmise that Gro is phosphorylated and, consequently, inactive at M-, G1- and G2-phases.

**Fig. 2.**
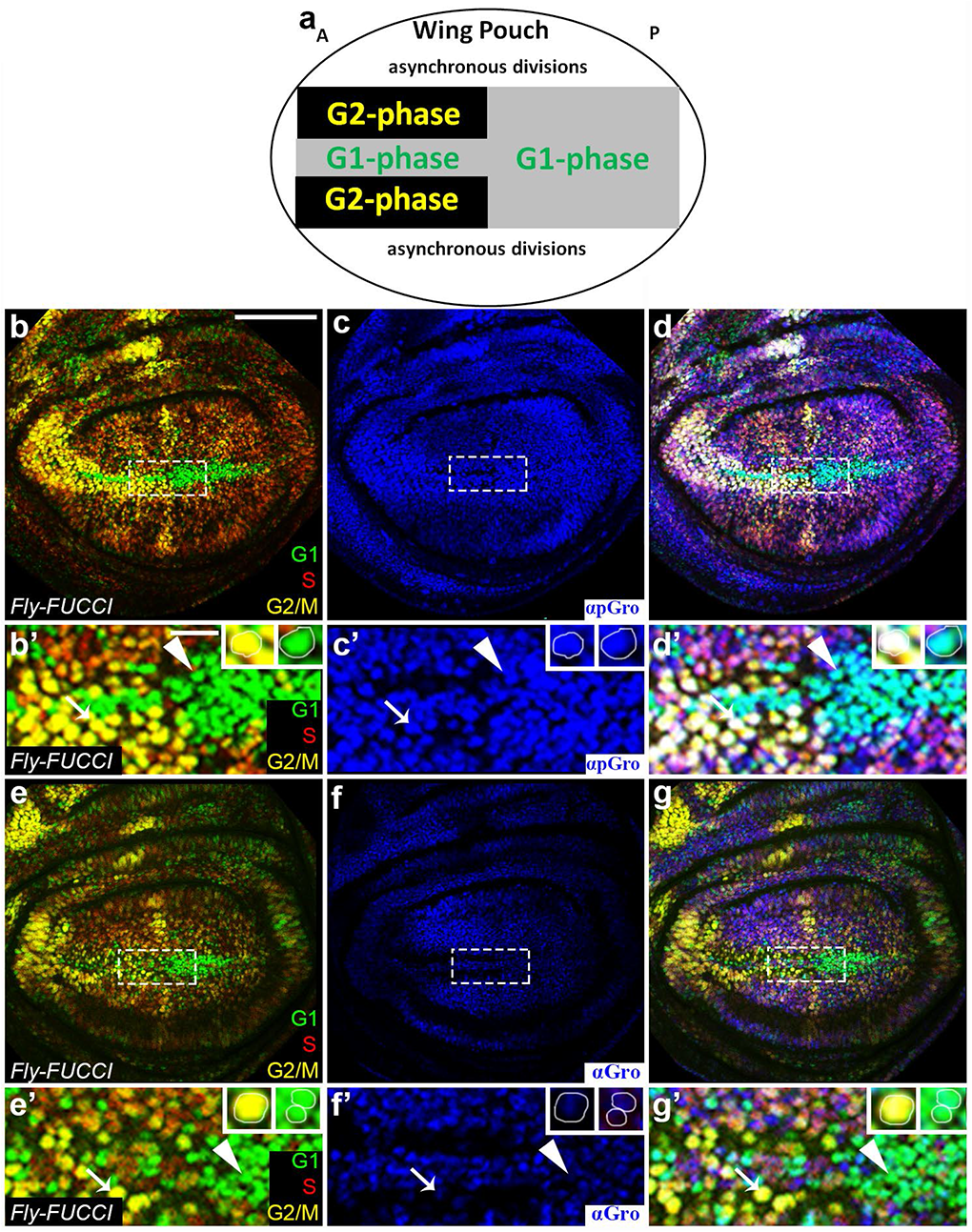
Groucho is phosphorylated during G1-, G2- and M-phases. (a) Schematic representation of the central part of the wing imaginal disc. In this region, a stripe of anterior cells that are arrested at G1-phase (grey) are flanked by cells arrested at G2-phase (black). Cells in the posterior domain are arrested at G1-phase [adapted from (Zielke et al., 2014)]. (b-g’) Confocal images of *Fly-FUCCI* third instar wandering larval wing imaginal discs, stained for pGro (blue; c-d’) or for Gro (blue; f-g’). In this system, the S-phase cell population is in red, cells at G2/M-phases are stained yellow and those at G1-phase are in green (b-b’, d-d’, e-e’, g-g’) (Zielke et al., 2014). (b’, c’, d’, e’, f’, g’) Magnified views of the central region of the wing imaginal disc blade shown in (b, c, d, e, f, g), respectively. Insets in panels of magnified views show representative cells either in G2/M (yellow; left) or in G1 (green; right). Note that pGro staining is evident in G2/M-phase nuclei, as well as in cells at G1-phase (arrow and arrowhead, respectively, in b’, c’, d’), but that of Gro is undetecatable in these cells (arrowhead and arrow in e’, f’, g’). The red, S-phase, marker alone was not analyzed due to intensity ambiguity. Scale bars = 100 µm (b, c, d, e, f, g) and 16.6 µm (b’, c’, d’, e’, f’, g’).

### Groucho is unphosphorylated, and therefore an active repressor, at S-phase

The *Fly-FUCCI* system also indicated that Gro is unphosphorylated at S-phase (Fig. 2). To confirm this, we carried out two additional experiments. First, wing imaginal discs of *PCNA-GFP* flies, in which expression of cytoplasmic GFP is a reliable marker for early S-phase (Strzalka and Ziemienowicz, 2011; Thacker et al., 2003), were stained for pGro. As shown in Fig. 3a-c, the pattern of GFP complements that of pGro (Fig. 3a-c). Second, we stained wing imaginal discs with the nucleoside thymidine analog 5-ethynyl-2’-deoxyuridine (EdU), which labels cells at late S-phase (Buck et al., 2008). Here, too, we find that nearly all EdU-positive cells co-stain for unphosphorylated Gro but not for pGro (Fig. 3d-f; quantified in Fig. 3j).

**Fig. 3.**
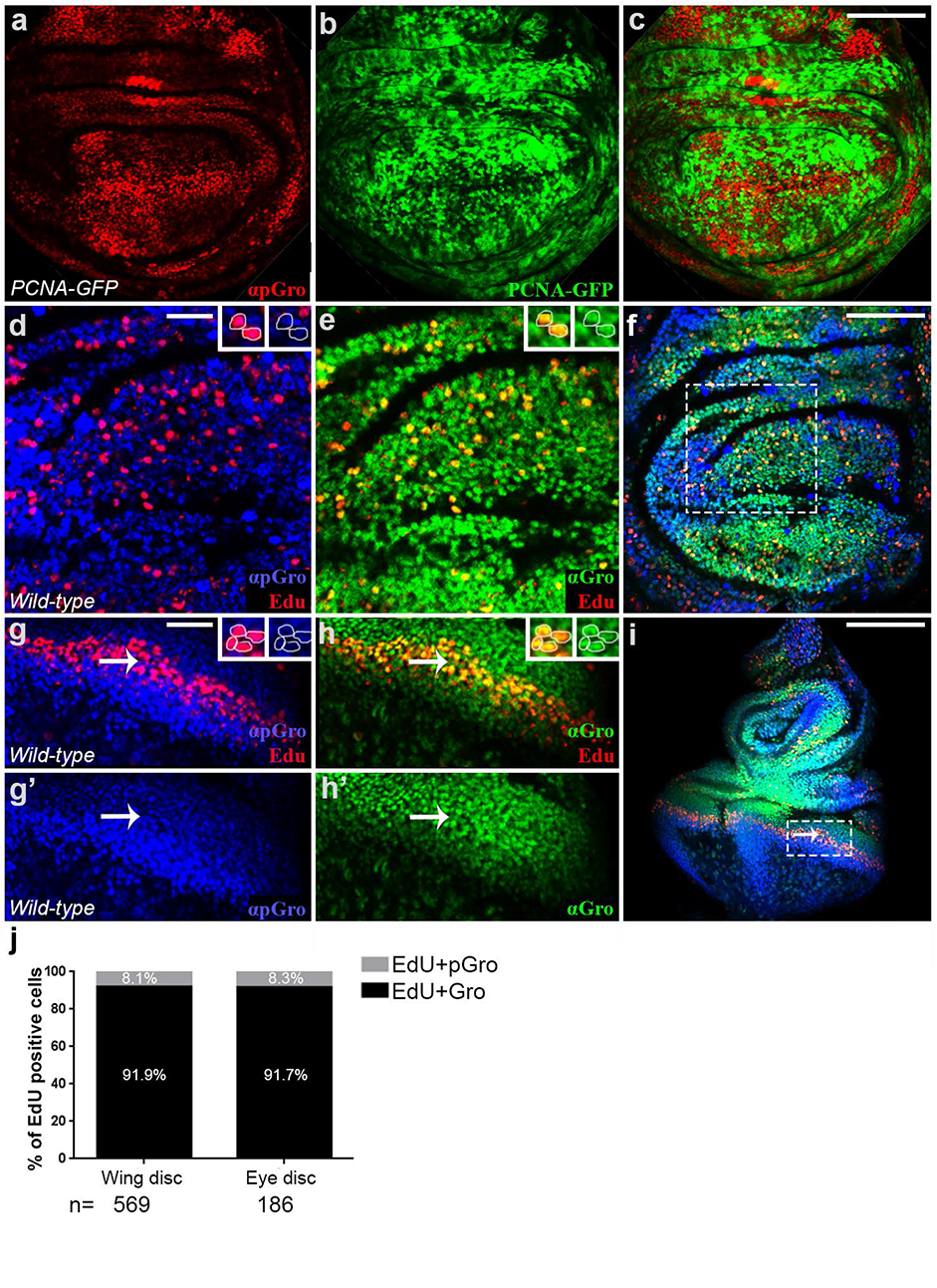
Groucho is unphosphorylated during S-phase. (a-c) Confocal image of *PCNA-GFP* third instar wandering larval wing imaginal disc, stained for pGro (red; a, c). GFP-positive, S-phase cells (green; b-c) do not stain for pGro. (d-i) Confocal images of third instar *wild-type* wandering larval wing (d-f) and eye (g-i) imaginal discs, stained for pGro (blue; d, f, g-g’, i), Gro (green; e, f, h-h’, i) and EdU (red; d-i). (d-e and g-h’) Magnified views of the boxed regions in (f and i), respectively. Arrow in (g-i) points at the stripe of EdU-positive, S-phase cells posterior to the morphogenetic furrow. Insets in (d-e, g-h) show magnified views of individual cells stained either for pGro and EdU (d, g) or for Gro and EdU (e, h). Note the overlap between EdU and anti-Gro staining, and the lack of correspondence between EdU and anti-pGro staining, in both wing and eye imaginal discs. (g) Percentage of EdU-positive nuclei co-stained with the anti-Gro antibody (black) or with the anti-pGro antibodies (grey) in wing and eye imaginal discs. n = number of EdU-positive cells scored in each case. Scale bars = 100 µm (a-c, f), 33.3 µm (d, e, g-h’) and 200 µm (i).

A similar result is also observed in eye imaginal discs, which allow the analysis of a relatively synchronized S-phase cell population. In this tissue, differentiation proceeds as a wave across the disc such that a stereotypic stripe of undifferentiated cells, located posteriorly to the morphogenetic furrow, synchronously enter S-phase (arrows in Fig. 3g-i) (Wolff and Ready, 1991). The vast majority of these EdU-positive S-phase cells stain for Gro but not for pGro (Fig. 3g-i; quantified in Fig. 3j), leading us to conclude that, in cycling cells, Gro is unphosphorylated only at S-phase, thereby restricting its repressive activity to this single stage of the cell cycle.

### Ectopic expression of Groucho reduces the number of mitotic cells

The non-overlapping patterns of anti-pH3 and anti-Gro staining (Fig. 1) suggests that Gro is phosphorylated in mitotic cells, and therefore inactive at this stage. To determine the significance of Gro phosphorylation to mitosis, we assessed the effects of expressing a non-phosphorylatable Gro mutant on the number of mitotic cells in the rapidly dividing wing imaginal disc. Towards this end, we employed a non-phosphorylatable Gro variant mutated in its two phosphoacceptor sites (Gro^AA^), in addition to a phosphomimetic Gro mutant derivative (Gro^DD^) (Cinnamon et al., 2008; Hasson et al., 2005; Helman et al., 2011; Helman et al., 2012; Johnston et al., 2016). If phosphorylation of Gro is a precondition for mitosis, we expect that expression of the constitutively active Gro^AA^ repressor should dominantly reduce the pH3 signal, but not that of Gro^DD^ whose repressor activity is attenuated.

The two variants were ectopically expressed in wing imaginal discs under Gal4/UAS control (Brand and Perrimon, 1993). Expression of Gro^AA^ using an early driver (*nubbin-Gal4*) resulted in small and dysmorphic discs, preventing their subsequent analysis. Instead, we utilized *MS1096-Gal4*, which drives non-uniform expression relatively late and mainly in the dorsal wing compartment (Suppl. Fig. S2). In this case, expression of Gro^AA^ but not of the converse phosphomimetic Gro^DD^ variant significantly reduces the number of pH3-positive cells relatively to control, LacZ-expressing wings (Fig. 4a-d, g-h), and the ensuing adult wings are markedly smaller (Suppl. Fig. S2). We conclude that phosphorylation of Gro is required for mitosis. In further support of this idea, there is a significantly greater overlay between the pH3 and Gro^DD^ signals (42% of 381 pH3-positive cells scored) than between those of pH3 and Gro^AA^ (14% of 254 pH3-positive cells scored) (Suppl. Fig. S2), indicating that cells expressing Gro^DD^, but not those expressing Gro^AA^, enter mitosis readily and that, therefore, phosphorylated Gro is compatible with M-phase.

**Fig. 4.**
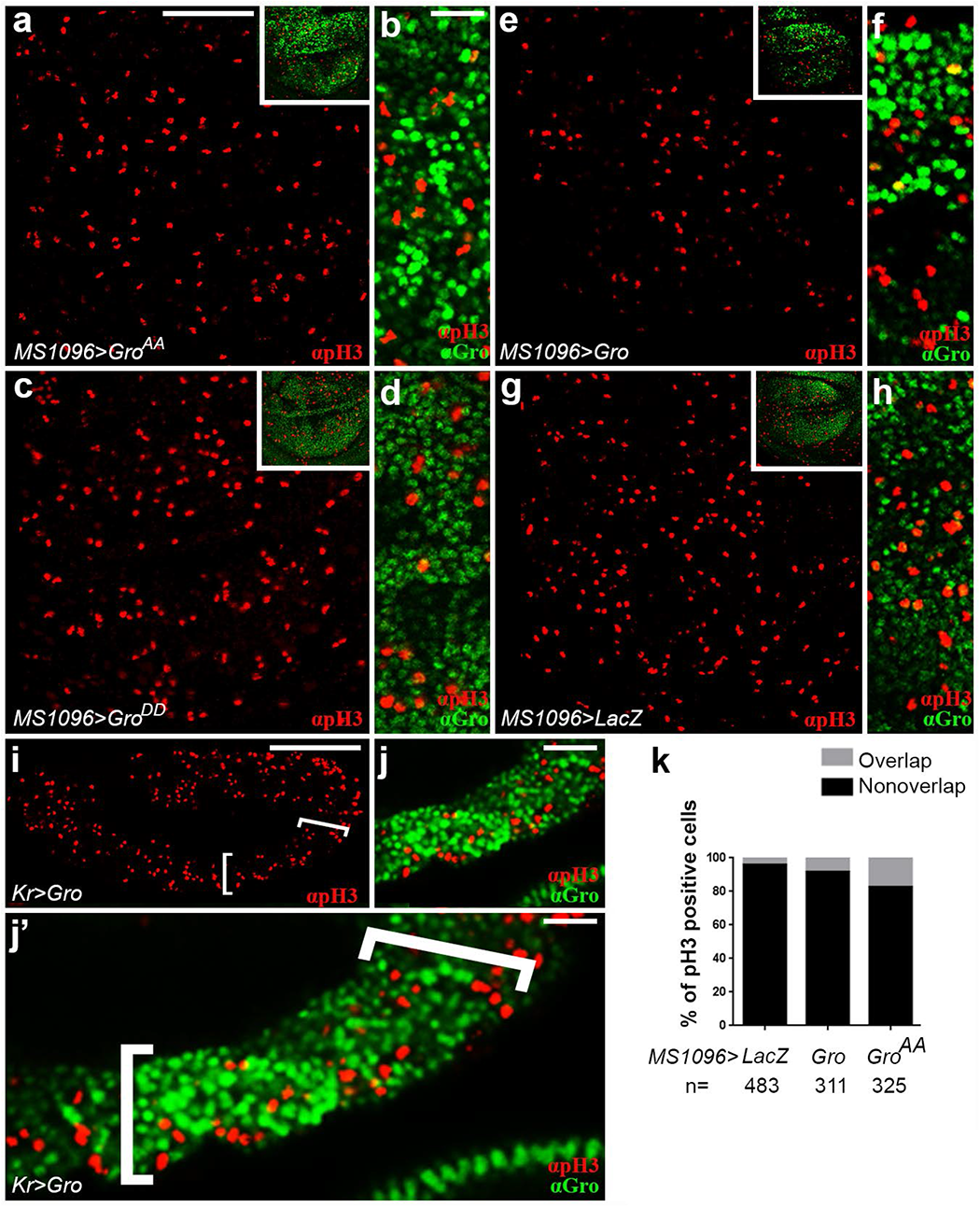
Ectopic expression of Groucho reduces the number of pH3-positive cells. (a-j’) Confocal images of wing imaginal discs (a-h) and stage 11 embryo (lateral view; i-j’), ectopically expressing a non-phosphorylatable Gro variant (Gro^AA^; a-b), a phosphomimetic Gro derivative (Gro^DD^; c-d) or native Gro (e-f, i-j’). LacZ-expressing wing disc served as control (g-h). Embryos and imaginal discs were co-stained for pH3 (red; a-j’) and for Gro (green; b, d, f, h, j-j’). Note that overexpression of Gro^AA^ and Gro in wing imaginal discs (a-b and e-f, respectively) decreases the number of pH3-positive nuclei relatively to Gro^DD^-expressing discs or to those expressing LacZ (c-d and g-h, respectively). Similarly, ectopic expression of Gro under *Kr>Gal4* regulation in the domain delineated by brackets (i-j’) reduces the number of pH3-positive nuclei. (b, d, f, h, j, j’) Magnified views of cells in panels (a, c, e, g and i), respectively. Note that the majority of persistent pH3-positive cells are those that essentially express minimal or null levels of Gro^AA^ or Gro. Insets in (a, c, e, g) show that ectopic expression of either Gro^AA^ (a) or Gro (e) masks the detection of endogenous Gro by the anti-Gro antibody, and that this anti-Gro antibody does not recognize ectopically-expressed Gro^DD^ due to its specificity towards unphosphorylated Gro (c). Hence, endogenous Gro is only observed in discs expressing Gro^DD^ or LacZ (c and g, respectively; see Materials and Methods). (b, e, j-j’) Patchy *Gal4*-driven expression leads to uneven Gro protein levels (Suppl. Fig. S3) (Barth et al., 2012; Ward et al., 2002). (k) Percentage of pH3-positive nuclei coinciding (grey) or not (black) with Gro staining in the indicated wing imaginal discs. n = number of pH3-positive cells scored in each case. Scale bars = 100 µm (a, c, e, g), 16.6 µm (b, d, f, h), 100 µm (i), 50 µm (j), and 33.3 µm (j’).

Induced expression of native Gro also causes a significant reduction in the number of pH3-positive mitotic cells, both in wing imaginal discs (Fig. 4e-f; *cf.* with control, LacZ-expressing disc in Fig. 4g-h) as well as in the embryo, which represent a different developmental tissue (using the *Krüppel* (*Kr*)-*Gal4* driver; Fig. 4i-j’) (Chu et al., 1998). To quantify Gro’s effects on mitosis, while controlling for changes its expression exerts on cell size, we calculated the ratio between the number of pH3-positive cells and the total number of cells in the wing pouch region (mitotic index). We find that the mitotic indices of wing imaginal discs overexpressing Gro (37.5%) and Gro^AA^ (46.8%), but not Gro^DD^ (87%), are significantly lower than the mitotic index observed in control, LacZ-expressing discs (related to as 100%). Notably, expression of native transcriptional regulators and their non-phosphorylatable derivatives, besides Gro and Gro^AA^, often exerts similar effects in other biological settings [see Fig. 4 in (Cinnamon et al., 2008); Figs. 3d-f and 4c in (Li et al., 2016); Figs. 5A-B in (Lee et al., 2010)].

**Fig. 5.**
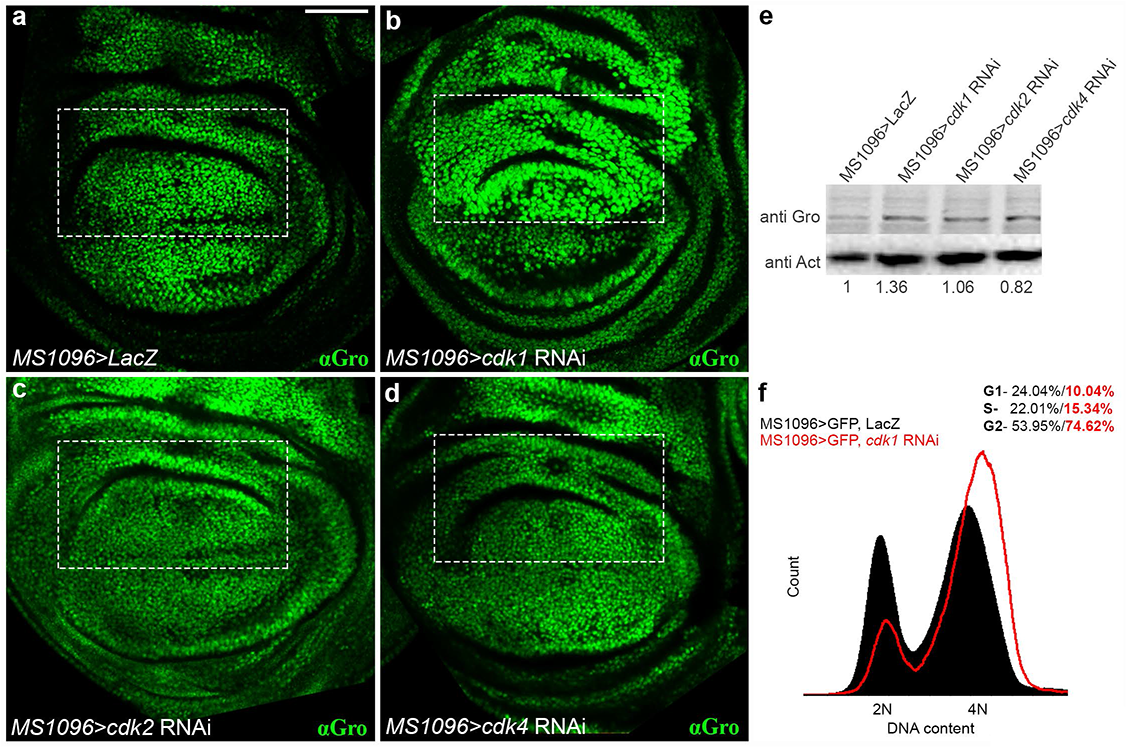
Cdk1 phosphorylates Groucho *in vivo*. (a-d) Confocal images of third instar wandering larval wing imaginal discs expressing either LacZ (a) or RNA interference (RNAi) constructs for *cdk1* (b), *cdk2* (c) or *cdk4* (d), stained for Gro (green; a-d). Two RNAi lines were used to target each Cdk, resulting in similar outcomes. The boxed regions in all panels demarcate the predominantly dorsal expression domain of the *MS1096-Gal4* driver (Suppl. Fig. S2). Note that *RNAi*-based knock-down of *ckd1* (b), but not of *ckd2* or *ckd4* (c and d, respectively), leads to the accumulation of unphosphorylated Gro (*cf.* control, LacZ-expressing disc in a). (e) Western blot analysis of whole wing imaginal disc lysates from the indicated genetic backgrounds, immunoblotted with anti-Gro and anti-Actin (anti-Act) antibodies. Relative Gro levels were determined based on the ratio between Gro and Actin, normalized to that in LacZ-expressing controls. The fold increase in the level of unphosphorylated Gro upon *cdk1* knockdown (2.222 ± 0.3975; *p*=0.0152; five biological repeats) is not observed in *cdk2* or *cdk4* knockdowns (1.152 ± 0.3023; *p*=0.51; and 1.045 ± 0.3030; *p*=0.8, respectively; three biological repeats for each; in all cases, data represents the mean ± SD, two-tailed *t*-test). Note that the increase in the level of unphosphorylated Gro following *cdk1* knockdown is probably a gross underestimate, given that *MS1096-Gal4* drives non-uniform expression in only a subset of cells in the wing imaginal disc (see Suppl. Fig. S2). (f) Cell cycle distribution of GFP-positive cells, dissociated from larval wing imaginal discs co-expressing either GFP together with LacZ (black) or along with *cdk1 RNAi* (red contour) under the *MS1096-Gal4* driver. DNA content was determined using Hoechst 33342 and normalized to number of events. Note the relative increase in the number of cells at G2/M-phases following *cdk1* knockdown. Scale bar = 100 µm (a-d).

Strikingly, irregular Gal4-driven expression (Suppl. Fig. S2) (Barth et al., 2012; Ward et al., 2002) revealed that the negative effect of Gro^AA^ and Gro on mitosis is largely cell-autonomous, since most of the remaining pH3-positive cells in the *MS1096-Gal4* and *Kr-Gal4* expression domains are consistently those that express null or low levels of induced Gro^AA^ and Gro (insets in Fig. 4b, h, f; quantified in Fig. 4k; Supplemental Movie 1). We conclude that entry into M-phase necessitates the prior attenuation of Gro’s repressive activity by phosphorylation.

### Cyclin-dependent kinase 1 phosphorylates Groucho *in vivo*

The above data indicate that phosphorylation of Groucho fluctuates dynamically in a cell cycle phase-dependent manner. Accordingly, Gro must be phosphorylated by a kinase that is active at G2-phase but inactive at S-phase. Although Erk activity impinges on cell cycle regulation at different levels (Mirzoyan et al., 2019; Mogila et al., 2006; Prober and Edgar, 2000), there is no evidence that it exhibits similar activation and inactivation dynamics, so Erk is probably not the kinase modulating Gro in this context.

Instead, we considered Cdk1 as the prime candidate for phosphorylating Gro (Vidwans and Su, 2001). Of the three fly Cdks, Cdk1 activity shows the appropriate dynamics for regulating Gro activity, being essential at G2- and M-phases but inactive at S-phase (Bettencourt-Dias et al., 2004). Additionally, Cdk1 (also known as cdc2) phosphorylates Gro and its human TLE1 orthologue *in vitro* and in cell culture (Nuthall et al., 2002). Finally, like Erk, Cdk1 is a Pro-directed Ser/Thr kinase (Malumbres, 2014) whose phosphorylation site in Gro was mapped to a fragment encompassing the Erk phosphorylation motif recognized by the pGro antibodies (see Materials and Methods) (Nuthall et al., 2002).

To test if Gro undergoes phosphorylation by Cdk1 *in vivo*, RNA interference (RNAi) was used to deplete it. Expression of *cdk1 RNAi* in wing imaginal discs results in fewer and larger cells, a known phenotype caused by Cdk1 deficiency (Suppl. Fig. S3) (Bettencourt-Dias et al., 2004) (Johnston, 1998). As Fig. 5 shows, the amount of unphosphorylated Gro increases in such *MS1096*>*cdk1 RNAi* wing imaginal discs, predominantly in the dorsal compartment (Fig. 5). The observed increase in unphosphorylated Gro is not simply due to a G2/M-phase arrest caused by *cdk1* depletion; *cdk1* knock-down leads to accumulation of cells at G2/M-phases (Fig. 5f) (Bettencourt-Dias et al., 2004), which are stages of the cell cycle in which Gro is normally phosphorylated (Fig. 2). Our results, therefore, probably reflect a direct role for Cdk1 in Gro phosphorylation *in vivo*, and are consistent with Cdk1 being the kinase responsible for inactivating Gro during G2 in this tissue.

### Phosphorylation of Groucho is mandatory for progression into mitosis

The Cdk1-mediated phosphorylation of Gro at the exit from S-phase appears to be obligatory for cell cycle progression. When Gro is overexpressed in *Fly-FUCCI* wing imaginal discs, a substantial enrichment in yellow-stained, G2/M-phase cells relative to control discs occurs (Fig. 6a-b; ImageJ measurements reveal a three-fold relative increase in the yellow-stained wing disc area) (Zielke et al., 2014). Since similar Gro expression leads to a cell-autonomous reduction in pH3 staining (Fig. 4), we conclude that the yellow staining reflects a significant increase in the number of cells at G2-phase.

**Fig. 6.**
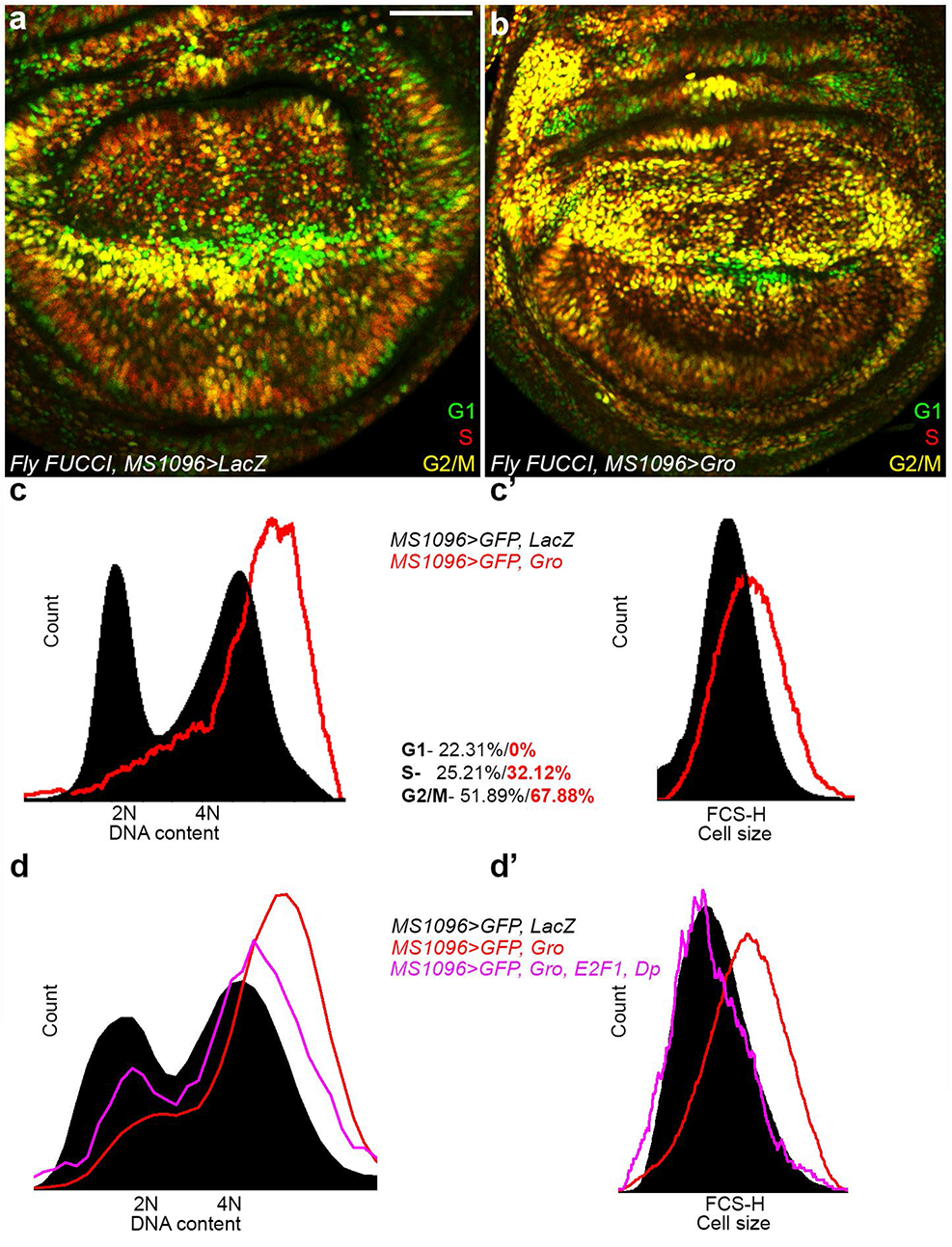
Ectopic expression of Groucho arrests cells at G2-phase. (a-b) Confocal images of wing imaginal discs, dissected from *Fly-FUCCI* third instar wandering larvae expressing either LacZ (a) or Gro (b) under *MS1096-Gal4* regulation. In the *Fly-FUCCI* background, cells at S-phase are stained red, those at G2/M-phases are stained yellow and the G1-phase population is in green (Zielke et al., 2014). Note that ectopic Gro expression significantly enriches the number of yellow, G2/M-phase cells in the dorsal compartment. (c-c’) Flow cytometric analyses of GFP-positive cells, dissociated from wing imaginal discs of flies expressing GFP together with LacZ (black), or GFP along with Gro (red contour), under the regulation of the *MS1096-Gal4* driver. The DNA content was determined using Hoechst 33342, and normalized to number of events. (c) Cell cycle distribution of GFP-positive, LacZ-expressing cells or of GFP-positive, Gro-expressing cells is depicted as percentages in black and red, respectively. Note the increased number of cells at G2/M-phases following Gro misexpression. (c’) Forward scatter-height (FSC-H) from the same experiment, showing cell size distribution. The relative size of cells from the Gro-expressing population (red contour) is generally larger than that of cells from the control population (black). (d-d’) Cell cycle distribution (d) and FSC-H reflecting cell size (d’) of GFP-positive cells, dissociated from larval wing imaginal discs co-expressing either GFP together with LacZ (black); GFP together with Gro alone (red contour); or GFP together with Gro, E2F1 and Dp (pink contour) under the *MS1096-Gal4* driver. DNA content was determined using Hoechst 33342 and normalized to number of events. Note that the concomitant expression of E2F1 and Dp partially suppresses the Gro-induced G2-phase arrest phenotype, as reflected by both cell cycle distributions and by cell size. Scale bar = 100 µm (a-b).

To confirm this result, we conducted flow cytometric analysis of dissociated wing imaginal disc cells. As Fig. 4c shows, a higher proportion of cells ectopically expressing Gro are in G2/M-phases compared to cells from control wings (Fig. 4c). Gro-expressing cells are also larger in size than control cells (Fig. 4c’) (Bettencourt-Dias et al., 2004). These results, together with the overlapping expression patterns of pH3 and pGro (but not Gro) staining (Fig. 1) and the Gro-induced decline in pH3-positive mitotic cells (Fig. 4), indicate that cells overexpressing Gro arrest at G2-phase, prior to mitosis, leading us to conclude that unphosphorylated, repressive Gro obstructs cell cycle progression.

### Groucho negatively regulates *e2f1* expression

An *in silico* approach was used to identify potential cell cycle targets of Gro-mediated repression. Multiple independent studies, using different methodologies, profiled Gro binding to chromatin in both *Drosophila* embryos and cultured cells (Suppl. Fig. S4) (Chambers et al., 2017; Kaul et al., 2014; Orian et al., 2007; Roy et al., 2010; Shibutani et al., 2008). Cell cycle and/or cell proliferation genes, closely located to Gro peaks, represent candidate Gro-regulated targets at S-phase. Moreover, the untimely transcriptional repression of some of these genes could also account for the G2-phase arrest imposed by overexpressed Gro (see Discussion).

From this gene set we chose, as case in point, the *e2f1* gene for further investigation. *e2f1* stands out as an attractive Gro target, since it encodes a key cell cycle regulator that activates genes required for the initiation of S-phase as well as for G2- and M-phases (Bennett et al., 1996; Dimova et al., 2003; Erez et al., 2008; Herr et al., 2012; Hurford et al., 1997; Ishida et al., 2001; Liu et al., 1996; Polager et al., 2002; Ren et al., 2002; Shibutani et al., 2008). The E2F1 protein is detectable at all phases of the cell cycle except for S-phase, when it is degraded (Bennett et al., 1996; Davidson and Duronio, 2012; Dimova et al., 2003; Herr et al., 2012). The inappropriate presence of E2F1 at S-phase can lead to accelareted cell cycle and/or apoptosis (Bennett et al., 1996; Davidson and Duronio, 2012; Dimova et al., 2003; Herr et al., 2012). While post-translational regulation of E2F1 has been extensively studied, regulation of *e2f1* gene expression is, as yet, largely unexplored (Goulev et al., 2008; Johnson et al., 1994; Øvrebø et al., 2022).

Several analyses support the notion that Gro represses *e2f1* gene expression during S-phase, and that phosphorylation of Gro attenuates this repression. Immunofluorescence staining of *wild-type* wing imaginal discs shows that signals of E2F1 and non-phosphorylated, repressive Gro do not overlap (Fig. 6a-a’) (Shibutani et al., 2008), and the pattern of the *PCNA-GFP* reporter, commonly used as proxy for E2F1 transcriptional activity (Buttitta et al., 2010), complements that of pGro (Fig. 3a-c). Moreover, ectopic non-phosphorylatable Gro^AA^ expression reduces anti-E2F1 staining, whereas Gro^DD^ expression does not affect E2F1 levels or distribution (Suppl. Fig. S4). Real-time polymerase chain reaction (RT-PCR) analysis reveals that overexpression of Gro in wing imaginal discs decreases the relative transcript levels of *e2f1* and several of its targets (Fig. 7b). Also, E2F1 is derepressed in homozygous *gro^E48^*mutant clones, induced in either wing (Fig. 7c-e) or eye imaginal discs (Fig. 7f-h). Finally, Gro is a direct or indirect repressor of *e2f1* transcription, rather than a controller of E2F1 post-translational levels, since it negatively regulates reporter expression derived from an *e2f1-lacZ* enhancer-trap line in eye imaginal discs (Fig. 7i-j) (Brook et al., 1996; Duronio et al., 1995).

**Fig. 7.**
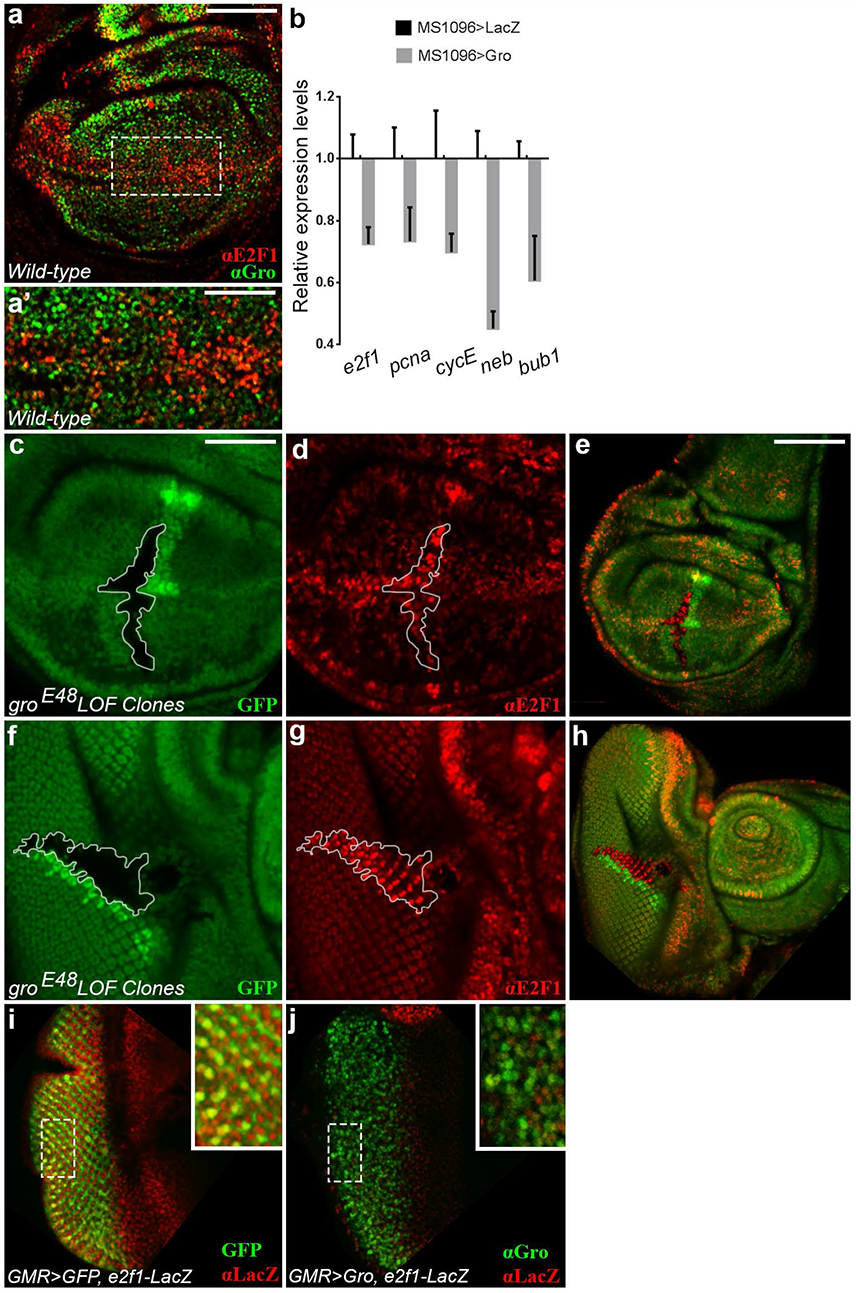
Groucho represses *e2f1* expression. (a-a’) Confocal image of *wild-type* third instar wandering larval wing imaginal disc, co-stained for E2F1 (red) and Gro (green). (a’) Magnified view of boxed region in (a). There is clear mismatch between the Gro and E2F1 nuclear signals. (b) RT-PCR analyses of mRNA extracted from third instar wandering larval wing imaginal discs expressing either Gro (grey) or LacZ (black) under *MS1096-Gal4* regulation. Relative transcript levels of *e2f1* and its G1- and G2-phase targets *pcna*, *cycE*, *neb* and *bub1* are reduced in Gro-expressing discs, normalized to LacZ controls. The ∼30% reduction in *e2f1* levels is probably an underestimation, given the mosaic expression driven by *MS1096>Gal4* (Suppl. Fig. S2). (c-h) Homozygous *gro^E48^ loss-of-function* clones (demarcated by white contours in c-d and f-g), discernable as GFP-negative and accompanied by GFP-positive twin spot clones (green; c, e, f, h), were induced in larval wing (c-e) and eye (f-h) imaginal discs. (c-d) and (f-g) are magnified views of clones in (e and h), respectively. E2F1 (red; d-e and g-h) is derepressed and ectopically accumulates in *gro* mutant clones. (i-j) Confocal images of third instar wandering larval eye imaginal discs, in which *GMR-Gal4* drives the expression of either GFP (green; i) or of Gro (green; j), co-stained for *e2f1-LacZ* reporter expression (LacZ; red; i-j). Insets show magnified views of the boxed regions in (i-j), respectively. (i) *GMR-Gal4* drives expression of GFP in differentiating retinal neurons (Yeates et al. 2019). These cells also express the LacZ reporter gene derived from the *e2f1-LacZ* enhancer trap. (j) Gro expression causes an overall reduction in anti-LacZ staining, particularly in the retinal neuronal cells overexpressing Gro (green), indicating that Gro negatively regulates *e2f1-LacZ* enhancer trap expression. Scale bars = 100 µm (a, c-d, i-j), 50 µm (a’, f-g) 200 µm (e, h).

We reasoned that if the G2-phase arrest induced by overexpression of Gro involves the inappropriate repression of *e2f1* at G2-phase, a stage when it should be re-transcribed, then the concomitant expression of E2F1 will rescue this phenotype. As Fig. 6 shows, the coexpression of E2F1 and Dimerization partner (Dp) suppresses the G2-phase arrest prompted by Gro overexpression (Fig. 6a-c’), and cell size reverts to normal (Fig. 6d-d’). This functional link between Gro and E2F1 reinforces the notion that relief of Groucho-mediated *e2f1* repression at G2-phase, via Cdk1-dependent phosphorylation, is crucial.

### Gro-mediated repression at S-phase is essential for proper cell cycle progression

The derepression of E2F1 does not induce apoptosis of *gro*-deficient cells, possibly because the level of upregulated E2F1 in *gro* mutant cells is still below the threshold required to trigger cell death, attainable by E2F1 overexpression (Suppl. Fig. S5) (Bennett et al., 1996; Davidson and Duronio, 2012; Dimova et al., 2003; Herr et al., 2012). To establish the significance of Gro-dependent repression at S-phase, we determined how genetic depletion of *gro* affects the fate of eye imaginal disc cells posterior to the morphogenetic furrow. These cells are normally in S-phase and, hence, are EdU-positive (Fig. 3g-i) (Wolff and Ready, 1991). When homozygous mutant *gro^E48^* clones intersect this domain, however, cells in which E2F1 is derepressed and consequently upregulated are consistently EdU-negative, indicating that they are not in S-phase (Fig. 8a-a’). The minority of *gro*-deficient cells that still stain for EdU, usually found at the margins of the clones, are typically E2F1-negative (Fig. 8a-a’; see Discussion).

**Fig. 8.**
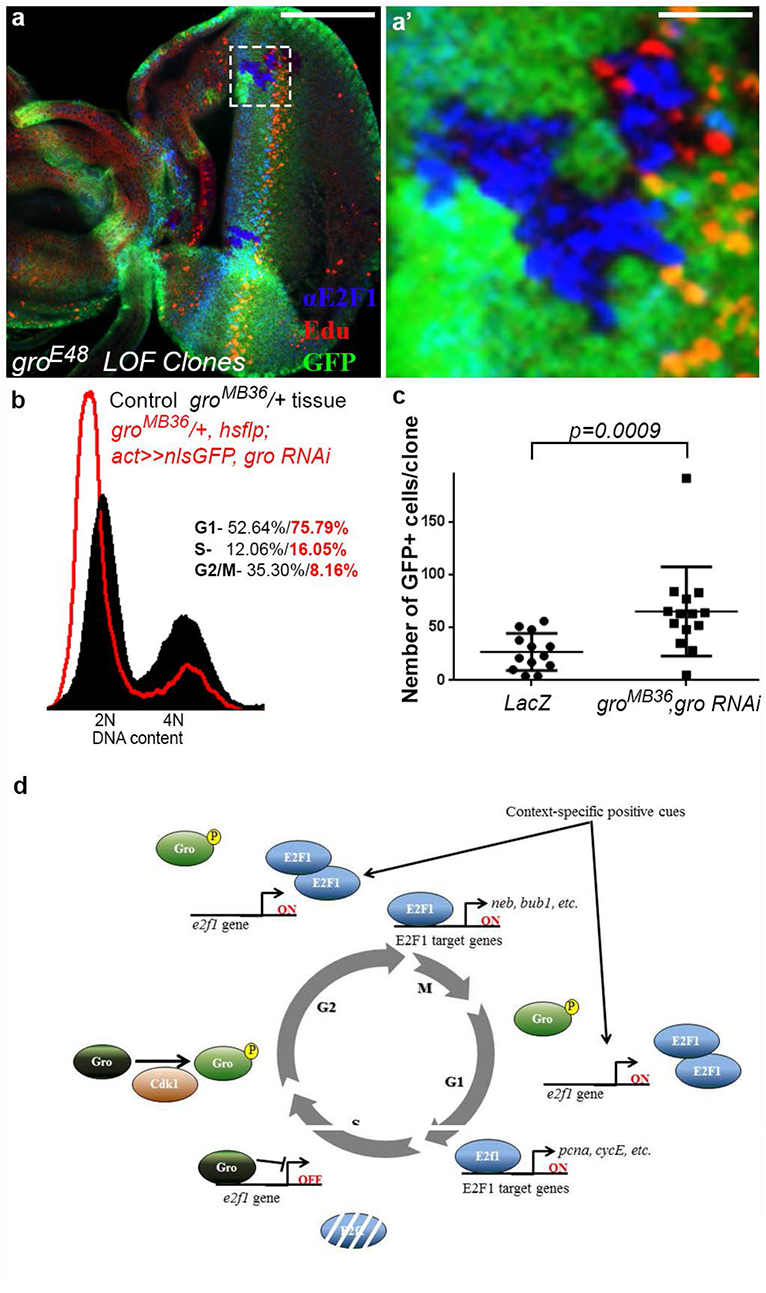
Cells devoid of *groucho* undergo accelerated cell cycles and accumulate at G1-phase. (a-a’) Confocal images of third instar wandering larval eye imaginal disc, stained for E2F1 (blue) and EdU (red). *gro* clones are detectable by lack of GFP staining and by their adjacent GFP-positive twin spot clone (green). (a’) Magnification of boxed region in (a), focusing on a *gro* mutant clone overlapping the morphogenetic furrow. Strikingly, all *gro* mutant cells that accumulate E2F1 do not stain for EdU and are, therefore, not in S-phase. (b) Flow cytometric analyses of dissociated cells from eye imaginal discs. *gro^MB36^*/+ cells expressing *gro RNAi* are labelled with GFP (red contour), whereas *gro^MB36^/+* cells that do not express *gro RNAi* are GFP-negative (black). The DNA content was determined using Hoechst 33342, and normalized to number of events. The cell cycle distribution of GFP-positive cells in which *gro* was downregulated, or of the remaining GFP-negative cells, is depicted as percentages in red and black, respectively. Note the increased number of cells at G1-phase following Gro downregulation. (c) The number of GFP-positive cells per clone, in *LacZ*-expressing (left) as well as in *gro^MB36^*/+ cells expressing *gro RNAi* (right) clones, under the regulation of *hsflp;actin>CD2>nlsGFP* driver. Note that RNAi-based reduction in Gro levels results in bigger clones, indicative of rapid cell cycles. 13 clones per each genotype were analyzed. *p*=0.0009 (Mann–Whitney *U*-test). (d) Schematic model. Please see text for details. Scale bars = 100 µm (a) and 16.6 µm (a’).

We hypothesized that if Gro-mediated repression at S-phase is critical, then perhaps its absence would perturb proper cell cycle progression. To address this point, we profiled the cell cycle distribution of GFP-marked eye clones of cells heterozygous for the *gro^MB36^*null allele, in which *gro* levels were further knocked-down (*gro RNAi*) (Suppl. Fig. S1) (Jennings et al., 2008). Flow cytometry analysis revealed that a significantly greater proportion of GFP-positive cells and, hence, with reduced *gro* levels have a 2N DNA content (Fig. 8b). Moreover, such eye imaginal disc clones show reduced staining for S-, G2- and M-phase markers (Suppl. Fig. S5). They, therefore, accumulate predominantly at G1-phase.

We considered two possibilities that can explain this result: *gro* depletion either i) leads to a cell cycle arrest prior to DNA synthesis or 2) accelerates the cell cycle such that cells pass through G2/M-phases faster. To distinguish between these options, we determined the doubling times of clonal cells overexpressing LacZ, or of cells in which *RNAi* was used to knockdown *gro*, by quantifying the number of GFP-positive cells per clone (Fig. 8c). Based on this data, we find that the respective doubling time of LacZ-expressing control clones is, on averge, 15.9h as previously reported (Tseng and Hariharan, 2002). Strikingly, *gro*-knockdown clones exhibit a significantly faster doubling time of 12h. The accelerated cell cycle phenotype of cells with lower levels of *gro* is remarkably similar to that caused by E2F1 overexpression (Davidson and Duronio, 2012; Shibutani et al., 2008), consistent with the idea that Gro normally represses *e2f1* at S-phase (Fig. 7). Moreover, the higher proportion of cells at G1-phase in *gro*-knockdown clones (Fig. 8b) resembles the accumulation of cells overexpressing E2F1 at G1-phase (Davidson and Duronio, 2012; Shibutani et al., 2008).

In summary, our data indicates that Gro repression is essential in S-phase for silencing *e2f1* expression and, possibly, of other genes (see Discussion). In Gro’s absence, the cycle accelerates and cells ultimately accumulate at G1-phase. Conversely, relief of Gro-mediated repression by Cdk1 activity is required for the correct progression through the G2-phase and for entry into mitosis (Fig. 8d). We conclude that Gro is an integral component of the cell cycle control system.

## Discussion

Our results uncover a previously unrecognized layer of cell cycle regulation by the Gro corepressor, indicating that Gro’s phosphorylation and dephosphorylation fulfil an integral regulatory function that is essential for cell cycle progression (Fig. 8d). We identify *e2f1* as a key target for repression by Gro at S-phase. Thus, Gro is a negative regulator of E2F1, a transcription factor that functions at the heart of the cell cycle by activating multiple genes required for the initiation of the cycle, as well as for the G2- and M-phases (Bennett et al., 1996; Dimova et al., 2003; Erez et al., 2008; Herr et al., 2012; Hurford et al., 1997; Ishida et al., 2001; Liu et al., 1996; Polager et al., 2002; Ren et al., 2002; Shibutani et al., 2008).

Previous reports had shown that preventing E2F1 proteolysis at S-phase brings about accelerated cell cycles and/or apoptosis (Davidson and Duronio, 2012; Shibutani et al., 2008). Repression of *e2f1* by Gro does not appear to provide a robust, backup mechanism for preventing E2F1-induced apoptosis, because *gro* mutant cells are viable despite the upregulation of E2F1 (Figs. 7, 8; Suppl. Fig. S5) (Hasson et al., 2005). We surmise that elevated E2F1 protein levels in *gro* deficient cells, in which the E2F1 degradation apparatus is presumably still operational, are not high enough to instigate cell death (Suppl. Fig. S5) (Davidson and Duronio, 2012). Instead, loss of Gro and the associated E2F1 upregulation leads to accelerated cell cycles.

Surprisingly, a minority of *gro*-deficient cells do not upregulate E2F1 and still stain for EdU (Fig. 8a-a’). It is conceivable that these EdU-positive cells, usually found at the periphery of *gro* clones, are exposed to non-cell autonomous signals from surrounding cells that drive them into S-phase, when E2F1 is robustly degraded. Their ability to continue dividing could explain why *gro* mutant clones are not eliminated.

Althought the negative regulation of *e2f1* could be indirect, due to repression by Gro of other genes that impinge on the cell cycle, several observations nevertheless point to a direct effect of Gro on *e2f1*. First, Gro binds to several chromatin sites within the first *e2f1* intron (Suppl. Fig. S4) (Chambers et al., 2017; Kaul et al., 2014; Orian et al., 2007; Roy et al., 2010; Shibutani et al., 2008), which is also included in the *e2f1*-enhancer trap line that is silenced by Gro (Fig. 7i-j) (Brook et al., 1996; Duronio et al., 1995). Second, E2F1 is repressed in cells arrested by ectopic Gro at G2-phase (Figs. 6, 7), despite the fact that it should be normally expressed at this stage (Davidson and Duronio, 2012; Shibutani et al., 2008). Third, Gro represses reporter gene expression derived from an *e2f1-lacZ* enhancer-trap line (Fig. 7i-j).

Surprisingly, Gro overexpression arrests cells at G2-phase, and not at G1-phase as expected from depletion of *e2f1* (Dobrowolski et al., 1994; Dyson, 1998; Fan and Bertino, 1997; Ishizaki et al., 1996; Wu et al., 1996). We surmise that overexpressed Gro blocks *de novo e2f1* transcription at G2-phase (following its repression by Gro at S-phase) and, consequently, Gro aborts progression of the cycle at this stage. In contrast, the G1-to-S-phase transition is refractory to Gro overexpression since the E2F1 protein has already accumulated, and is post-transcriptionally regulated (via Rb and its phosphorylation), at this stage of the cycle. Consistent with this interpretation, coexpression of E2F1 and Dp suppresses the G2-phase phenotypes induced by Gro overexpression (Fig. 6d-d’).

In our work we have focused on *e2f1* as a key target regulated by Gro. However, it is also conceivable that Gro represses additional genes, in proximity to which it binds, that must be silenced during S-phase. Prospective candidates for Gro repression are genes encoding the *auroraA* and *auroraB* kinases that are required for the control of mitosis (Goldenson and Crispino, 2015; Orian et al., 2007), and *transforming acidic coiled coil* (*TACC*) which maintains spindle bipolarity and microtubule stability during mitosis (Trivedi, 2013). We do not rule out the possibility that silencing of these (and perhaps other) genes contributes to the predominant G2-phase arrest exerted by Gro expression (Fig. 6).

The majority of pGro-positive cells are negative for pH3 staining and are, therefore, not mitotic (Fig. 1). These could be non-dividing cells or, if cycling, could be in one of the two Gap-phases at which Gro is also phosphorylated (Fig. 2). Regardless, we propose that for cell division to occur, the attenuation of Gro repression must be accompanied by specific pro-proliferative cues (Fig. 8d). Similar context-dependent positive inputs, each exclusive to a particular cellular process, will ensure that the downregulation of Gro by phosphorylation results in induction of a restricted sets of genes in each case, depending on the cellular context.

The full scope of Gro’s regulatory functions in the framework of the cell cycle will come to light once the complete repertoire of its targets is revealed and its DNA-binding partner proteins in this framework identified. How phosphorylated Gro is replaced by unphosphorylated Gro at S-phase also remains an open question. We assume that either Gro is dephosphorylated by a Ser/Thr phosphatase, or that phosphorylated Gro is degraded at G1- or early S-phase, with non-phosphorylated Gro newly transcribed and translated during S-phase. Further studies will be required to discern between these possibilities.

To date, Gro has been mainly implicated in transcriptional repression events controlling cell fate specification and differentiation (Cinnamon and Paroush, 2008; Hasson and Paroush, 2006). The new roles we have uncovered for Gro and its phosphorylation in the cell cycle raise the possibility that they also function in additional basic cellular processes. Other kinases, besides Erk and Cdk1, have been shown to phosphorylate Gro (Choi et al., 2005; Nuthall et al., 2002; Nuthall et al., 2004), and software programs predict yet additional kinases that could potentially modify the motif recognized by the anti-pGro antibodies. Accordingly, other physiological and/or metabolic processes, each employing a specific effector kinase(s), may derepress their unique arrays of downstream target genes via phosphorylation of Gro.

Emerging evidence supports the notion that Gro’s human TLE orthologues may act similarly to Gro in the context of the cell cycle. TLE1 and TLE3 are both implicated in cancer; TLE1, for example, promotes glioblastoma propagation (Dali et al., 2018) and TLE3 stimulates cell division by suppressing myogenic differentiation via transcriptional repression of the master regulator MyoD (Kokabu et al., 2017). Moreover, a large-scale analysis found that TLE1 is a mitotic bookmarking factor in development and in stem cells (Festuccia et al., 2017), and TLE3 was identified in a phosphoproteomics analysis of full phosphorylation site occupancy during mitosis (Olsen et al., 2010). Finding that TLE corepressors function comparably to Gro in cell cycle regulation may ultimately offer new therapeutic strategies for preventing uncontrolled cell division in cancerous settings.

## Materials and Methods

### Fly culture and stocks

Flies were cultured and crossed on standard yeast-cornmeal-molasses-malt extract-agar medium at 25°C.

cDNAs for Gro, Gro^AA^ and Gro^DD^ were cloned into the *pUAST-attB* vector, and all constructs were subsequently integrated at the *attp40* site (BestGene Inc., USA) to generate transgenic lines with comparable expression levels (Suppl. Fig. S2d) (Markstein et al., 2008).

The following GAL4 drivers and responders were used: *kr-Gal4* (BDSC #58800); *MS109*6-*Gal4*; *gro^MB36^*, *UAS-gro RNAi*/*TM6B* (generously provided by Gerardo Jiménez) (Jennings et al., 2008); *UAS-cdk1 RNAi* (BDSC #28368 and #36117); *UAS-cdk2 RNAi* (BDSC #28952 and #34856); *UAS-cdk4 RNAi* (BDSC #36060 and #27714); *UAS*-*LacZ* (Brand and Perrimon, 1994); *UAS-GFP* (Yeh et al., 1995); *UAS*-*E2F1, UAS*-*Dp/CyO* (BDSC #4774); *PCNA-GFP* (kind gift of Robert Duronio); *P[rm729] e2f1-lacZ* (BDSC #34054); and *Fly-FUCCI* (BDSC #55124 and #55123) (Zielke et al., 2014). *yellow white* flies served as *wild-type* controls.

### Generating *groucho* loss-of-function and overexpression clones

Mutant clones lacking functional Gro (*gro^E48^*) were generated using FLP-mediated mitotic recombination in the progeny of the following cross: *hsflp*; *P[FRT82B] ubi-GFP/TM6B* virgin females and *P[FRT82B] gro^E48^/TM6B* males. Clones were induced 48-72hr after egg laying by heat-shock (60min at 37°C), and were identified by the loss of the GFP marker and the concurrent appearance of a twin spot clone. Gro knock-down was attained by crossing *hsflp*; *actin>CD2>Gal4*; *UAS-nlsGFP/TM6B* virgins to *gro^MB36^*, *UAS-gro RNAi*/*TM6B* males (Jennings et al., 2008). Clones, induced 48-72hr after egg laying by heat-shock (10min at 37°C), were distinguishable via the GFP marker.

### Preparing wing imaginal disc lysates for immunoblotting

Wing imaginal disc lysates were prepared as described (Kushnir et al., 2020).

### Antibody staining

Primary antibodies used in this study were: rabbit anti-phospho-Histone 3 (1:100; Cell Signaling #9701); mouse anti-phospho-Histone 3 (1:100; Cell Signaling #9706); rabbit anti-pGro (1:100) (Cinnamon et al., 2008; Hasson et al., 2005); mouse anti-Gro (diluted 1:1000 for immunofluorescence and 1:5000 for Western blot analysis; generously contributed by Christos Delidakis) (Delidakis et al., 1991); rat anti-total Gro (1:1000 for Western blot analysis; Santa Cruz sc-15786); rabbit anti-panTLE (1:100, Cell Signaling #4681); mouse anti-GFP (1:100; Developmental Studies Hybridoma Bank (DSHB) #8H11); mouse anti-Cyclin A (1:20; DSHB #A12); mouse anti-Cyclin B (1:20; DSHB #F2F4); rabbit anti-Dcp1 (1:100; Cell Signaling #9578); and rat anti-E2F1 (1:100; generous gift by Stefan Thor). Secondary antibodies were 488-, Rhodamine Red-X- or Cy5-conjugated (1:400; Jackson Laboratories).

Nuclei were labeled using DAPI (1:1000; Sigma), and embryos and wing imaginal discs were mounted using Vectashield medium (Vector Laboratories, Inc.).

### Immunovisualisation of Gro’s phosphorylation state *in vivo*

Rabbit anti-phosphorylated-Gro (pGro) polyclonal antibodies were raised using a synthetic phosphopeptide containing one of two putative Erk consensus sites (phospho-Thr308). Subsequently, the antibodies were affinity-purified on a column with a corresponding non-phosphorylated peptide, and the flow-through was later bound on a column with the phosphorylated peptide. These anti-pGro antibodies detect Gro in its phosphorylated state *in vivo*, particularly in domains of ongoing and earlier (given the persistence of Gro phosphorylation) RTK pathway activity (Cinnamon et al., 2008; Hasson et al., 2005; Helman et al., 2011; Johnston et al., 2016).

Immunofluorescence staining, using a monoclonal mouse anti-Gro antibody raised against amino acids 120-380 (Delidakis et al., 1991), generates a signal that largely complements the domain of pGro throughout embryonic and adult development (Suppl. Fig. S1) [see Figs. 1-3 in (Cinnamon et al., 2008) and Fig. 5 in (Johnston et al., 2016)]. Note that the anti-Gro antibody detects signals across a range of Gro protein levels (i.e., downregulation as well as overexpression; Suppl. Fig. S1 and Fig. 4). The general lack of anti-Gro staining in anti-pGro-positive nuclei implies that the protein is completely phosphorylated in these nuclei. Importantly, the mutually exclusive recognition by the anti-pGro and anti-Gro antibodies is also observed *in vitro* [(Fig. 1 in (Cinnamon et al., 2008)], indicating that phosphorylation is enough to cause differential detection of the same, or neighboring, epitopes by the two anti-sera.

The specific recognition of nonphosphorylated Gro by the anti-Gro antibody is further illustrated by its failure to detect phosphomimetic Gro^DD^, while strongly recognizing the nonphosphorylatable Gro^AA^ variant when similarly overexpressed (*cf*. insets in Figs. 4c and 4a, respectively). Finally, commercially available polyclonal anti-total Gro antibodies (Santa Cruz) were used to detect total Gro levels (i.e., Gro, Gro^AA^ and Gro^DD^) in immunoblots (Suppl. Fig. S2), and commercially available anti-panTLE antibodies (Cell Signaling) were used to follow transgenic expression *in vivo* (Suppl. Fig. S2).

### EdU incorporation

Third instar larval wing and eye imaginal discs were submerged in 1xPBS in the presence of 1:1000 EdU for 1hr with gentle rolling at RT. EdU was detected using the Click-iT EdU Alexa Fluor 555 Imaging Kit (Life Technologies).

### Real-time polymerase chain reaction (RT-PCR)

Total RNA was extracted using Aurum Total RNA Mini Kit (Bio-Rad). RT-PCR was performed using iTaq Universal SYBR Green Master Supermix (BioRad) and CFX384 Touch Real-Time PCR Detection System (Bio-Rad). Expression was normalized to *GAPDH* transcripts in all cases, and each experiment was carried out in biological triplicates (one representative experiment is shown). Data analysis was preformed using Bio-Rad CFX manager 3.1 (Bio-Rad).

### Flow cytometry

Samples were prepared as described (Davidson and Duronio, 2012), and DNA content was determined using Hoechst 33342 (Sigma). Cells were subjected to flow cytometry using LSR-Fortessa Analyzer (BD Biosciences), and results were analyzed by FCS express 4 software (De-Novo software). Each experiment was carried out at least three independent times, of which a single representative experiment is presented.

### Quantifying mitotic indices and doubling time of cell populations

Mitotic indices, which take into account Gro’s effect on mitosis while controlling for the changes its expression exerts on cell size, were calculated by counting pH3-positive cells in a known, constant area and dividing them by the number of nuclei.

For doubling time measurement, clones were induced by heat shock at 72h and fixed at 124h. Doubling time was calculated using the following formula: (Log 2/Log N)hr, where N is the median cell number per clone and hr is the age of the clones in hours (Tseng and Hariharan, 2002).

### Mounting of fly wings

Adult wings were mounted as described in (Kushnir et al., 2020).

### Microscopy

Confocal images were attained using a Zeiss LSM710 confocal microscopy. Images were processed using Adobe Photoshop software.

### Statistical analyses

Statistical analyses were conducted using Prism software. ImageJ was used to measure the intensity of yellow-stained cells and Zen software to quantify the numbers of DAPI, pH3-, EdU-, Gro- or pGro-positive cells.

## Supplemental Information

**Supplemental Fig. S1.**
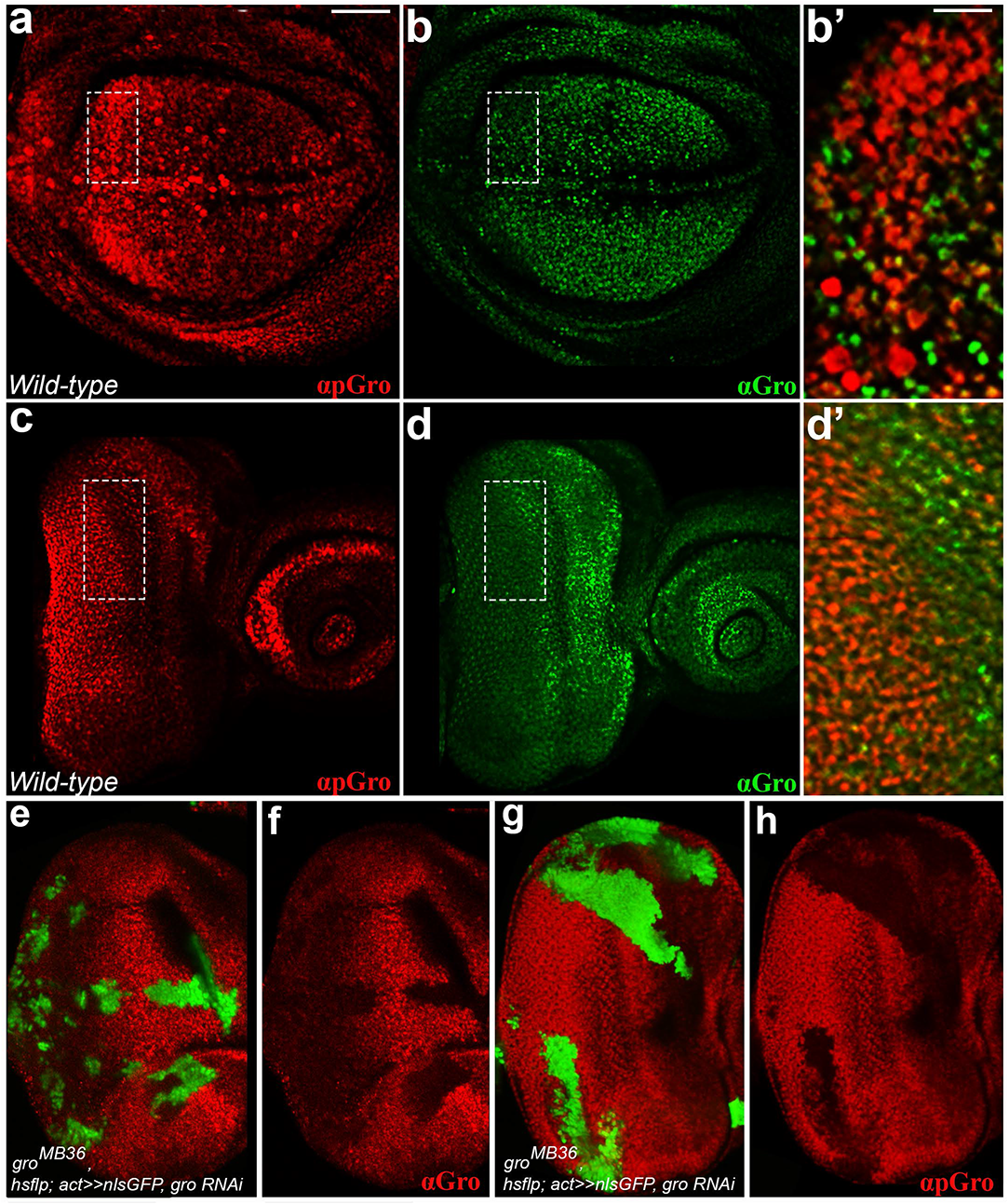
Immunovisualisation of Groucho’s phosphorylation state *in vivo* using anti-Gro and anti-phospho-Gro antibodies. (a-d’) Confocal images of *wild-type* third instar wandering larval wing (a-b’) and eye (c-d’) imaginal discs, co-stained for pGro (red; a, b’, c, d’) and Gro (green; b-b’, d-d’). (b’, d’) Magnified views of boxed regions in (a-b and c-d), respectively. (a-d’) The complementarity in epitope detection by the anti-pGro and anti-Gro antibodies is evident (Cinnamon et al., 2008; Johnston et al., 2016). The relative prevalence of anti-pGro staining compared to that of anti-Gro probably stems from the persistence of Gro phosphorylation (Helman et al., 2011). (e-h) Both the anti-Gro antibody and the anti-pGro antibodies are sensitive to RNAi-mediated reduction in Gro levels. Eye imaginal discs in which *gro* was knocked-down in clones of cells heterozygous for the *gro^MB36^* allele, discernable by GFP staining (green; e, g), were stained for Gro (red; e-f) or for pGro (red; g-h). Note that reduced intensity of the anti-Gro (e-f) and anti-pGro (g-h) signals in the clones. Scale bar = 100 µm (a-b, c-d, e-h) and 33.33 µm (b’, d’).

**Supplemental Fig. S2.**
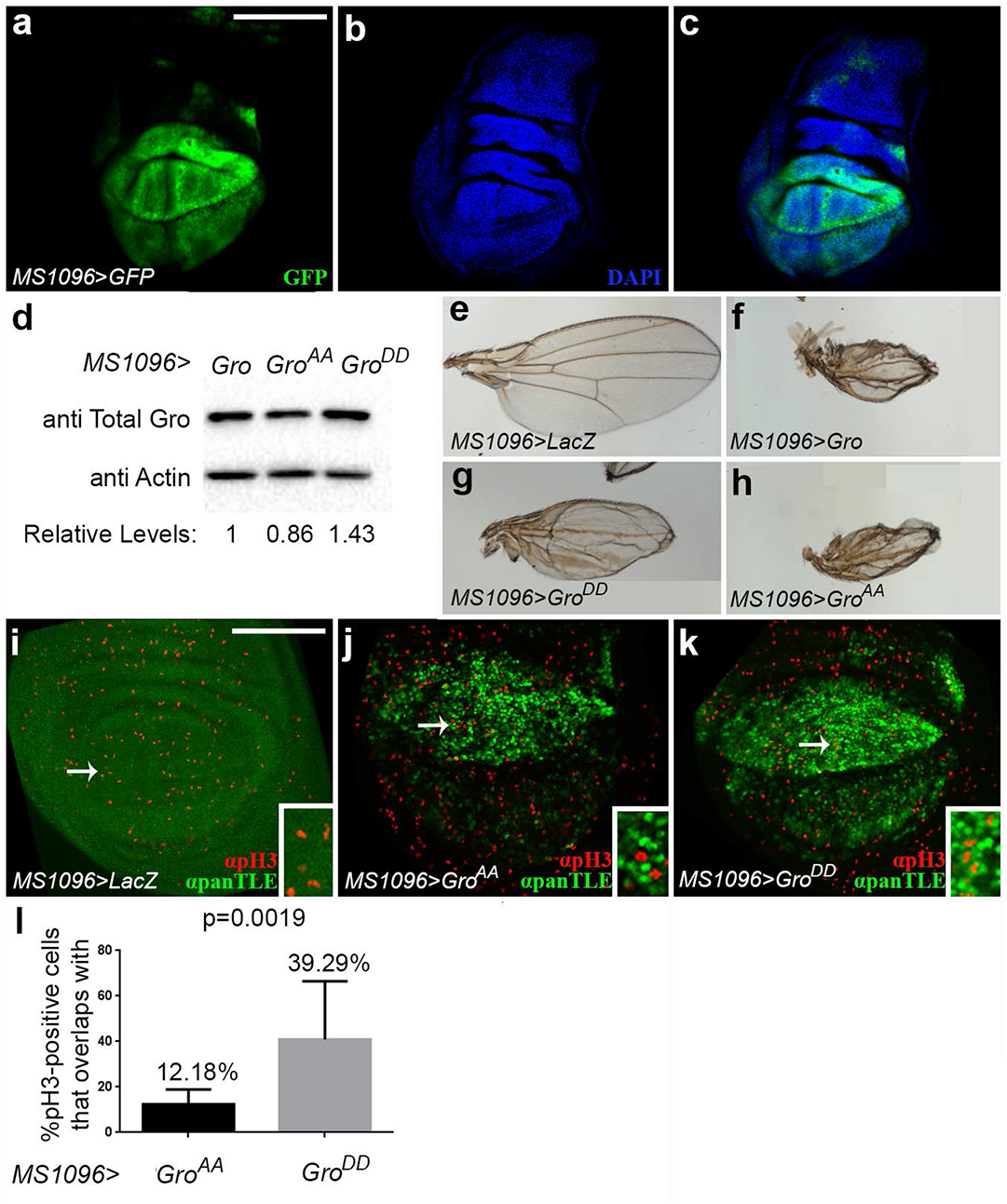
Transgenic expression of Gro, Gro^AA^ and Gro^DD^ using the *MS1096-Gal4* driver. (a-c) Confocal image of third instar wandering larval wing imaginal disc, in which *MS1096-Gal4* drives expression of GFP (green; a, c), counterstained for DAPI (blue; b-c). GFP is expressed predominantly in the dorsal region of the wing imaginal disc, but also in some cells in the ventral compartment (albeit to a lesser extent), probably due to leakiness of the *MS1096-Gal4* driver. Note the uneven, irregular nature of the UAS/Gal4 overexpression system. (d) Immunoblot analysis showing that relative transgenic expression levels of Gro, Gro^AA^ and Gro^DD^, driven by *MS1096-Gal4*, are comparable. Relative levels of Gro^AA^ and Gro^DD^, determined based on the ratio between total Gro and Actin levels, were normalized to these values for Gro. (e-h) Wings of adult females of the indicated phenotypes. (i-k) Confocal images of third instar wandering larval wing imaginal discs, overexpressing LacZ (i), Gro^AA^ (j) or Gro^DD^ (k) under the regulation of the *MS1096-Gal4* driver, co-stained for panTLE (green) and pH3 (red). Insets in (i-k) show magnified views of regions marked by respective arrows. Note that anti-panTLE antibodies, which are insensitive to Gro’s phosphorylation state and therefore recognize both Gro^AA^ and Gro^DD^, weakly detect endogenous Gro (i) but visibly detect overexpressed Gro^AA^ (j) and Gro^DD^ (k). (l) A significantly larger proportion of pH3-positive mitotic cells overlaps with Gro^DD^ than with Gro^AA^. Data represents the mean ± SD derived from 10 wings per genotype. (*p*=0.0019; Mann–Whitney *U*-test). Scale bars = 50 µm (a-c) and 100 µm (i-k).

**Supplemental Fig. S3.**
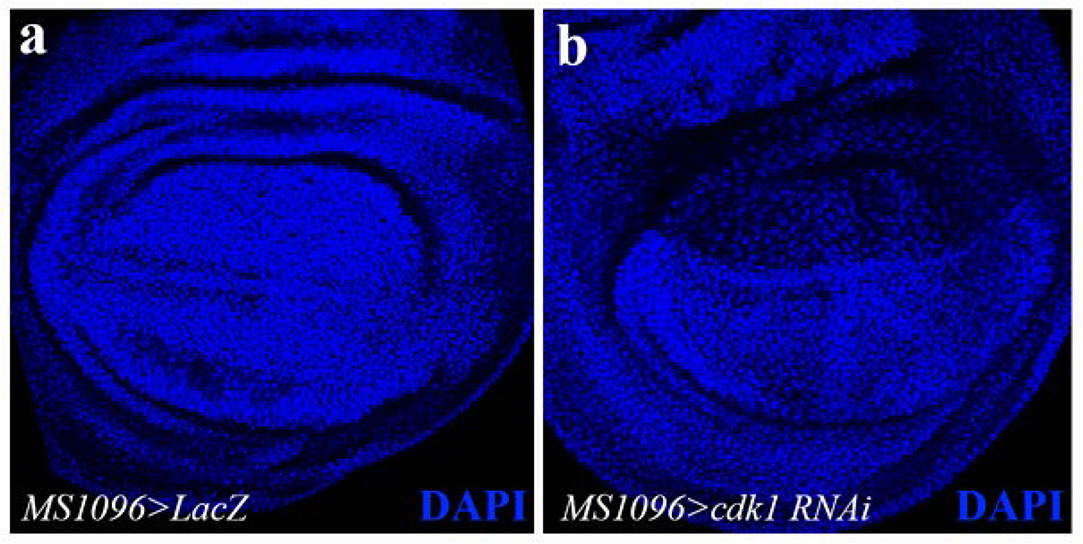
RNA interference-based reduction in Cdk1 levels results in fewer and larger cells. (a-b) Confocal images of third instar wandering larval wing imaginal discs, expressing either LacZ (a) or an RNA interference (RNAi) construct for *cdk1* (b), stained with 4’,6-diamidino-2-phenylindole (DAPI) (blue). Note that fewer and larger nuclei are observed in the domain of *cdk1* downregulation, as previously reported (Bettencourt-Dias et al., 2004); (Johnston, 1998). Scale bar = 100 µm.

**Supplemental Fig. S4.**
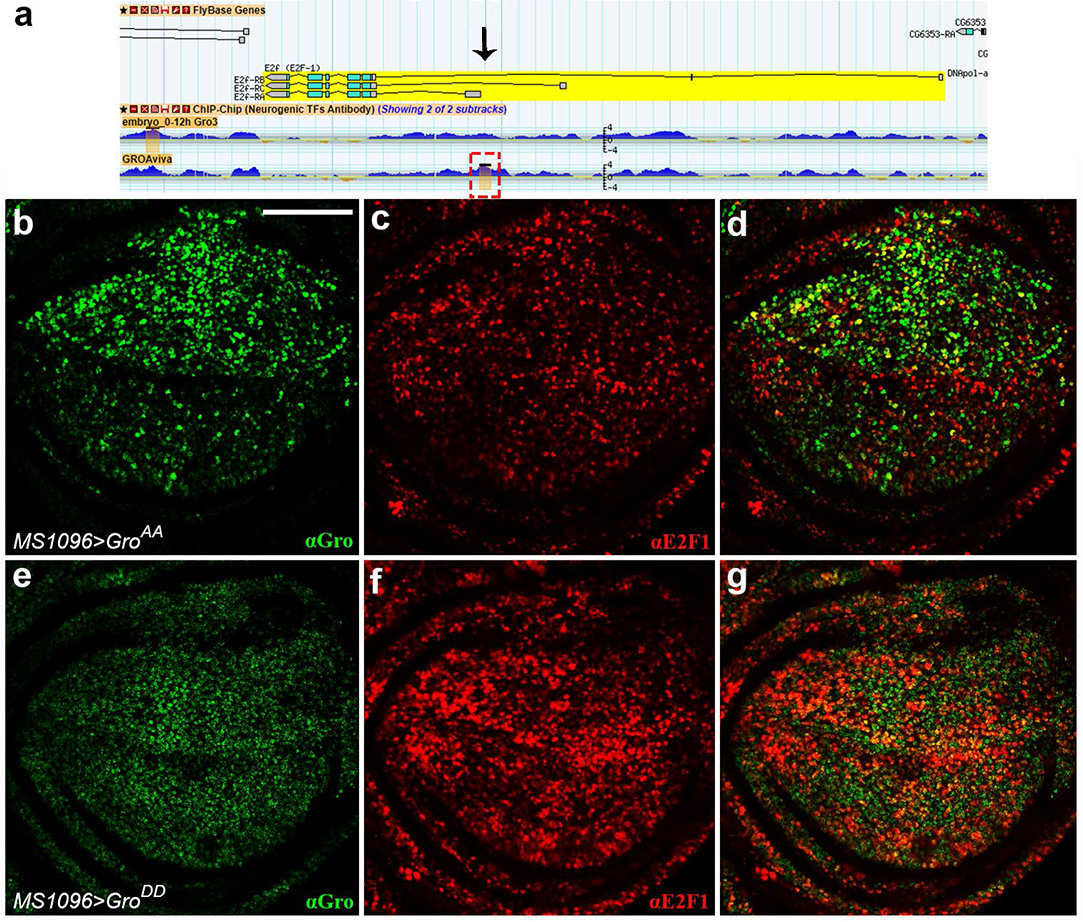
Gro^AA^, but not Gro^DD^, represses *e2f1* expression. (a) Gro binds ajacent to the *e2f1* locus. Shown is a schematic display from the modENCODE Project (http://www.modencode.org/) depicting significant Gro binding to a region within the first intron of the E2f-RB and E2f-RC transcripts and next to the transcription start site of transcript E2f-RA. Other independent studies also mapped Gro binding to this region. (b-g) Confocal images of third instar wandering larval wing imaginal discs overexpressing Gro^AA^ (b-d) or Gro^DD^ (e-g) under the regulation of the *MS1096-Gal4* driver, co-stained for Gro (green; b, d, e, g) and E2F1 (red; c-d, f-g). Note the overall decrease in anti-E2F1 staining in the Gro^AA^-expressing disc (c), in comparison to the disc expressing Gro^DD^ (f). Scale bar = 100 µm.

**Supplemental Fig. S5.**
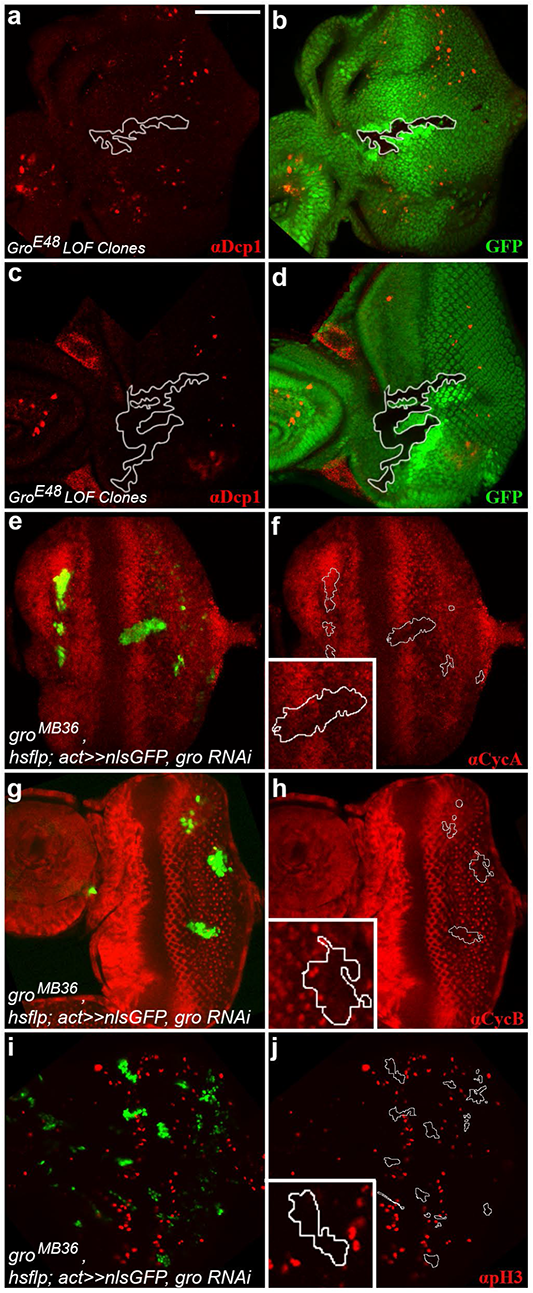
Cells with reduced Groucho levels do not undergo apoptosis, and do not stain for S-, G2- and M-phase markers. (a-d) GFP-negative (green; b, d) homozygous *gro^E48^* loss-of-function clones (demarcated by white contours), induced in larval eye imaginal discs, were stained for the activated form of the *Drosophila* effector caspase, *Drosophila* caspase 1 (Dcp-1) (red; a-d). Anti-Dcp-1 staining is not elevated in *gro* clones. (e-j) Confocal images of third instar wandering larval eye imaginal discs, in which *gro* was knocked-down in GFP-labelled clones of cells heterozygous for the *gro^MB36^* allele (green; e, g, i). (f, h, j) Clonal boundaries are outlined, with each inset showing a magnified view of a representative clone. Note that the majority of cells, in which *gro* levels are reduced, do not stain for the S- and G2-phase marker CycA (red; e-f); for the G2-phase marker CycB (red; g-h); or for the mitotic marker pH3 (red; i-j). Scale bar = 100 µm.

**Supplemental Movie 1.** Z-stack imaging of a stage 11 *Kr>Gro* embryo, co-stained for pH3 (red) and Gro (green). Note that the two signals are non-overlaping.

## Acknowledgments

We thank Tatyana Kushnir and Rotem Lange for continuous help and encouragement during this project. We are grateful to Einat Cinnamon, Russell Finley Jr., Offer Gerlitz, David Ish-Horowicz, Gerardo Jiménez, Michael Klutstein and Benny Shilo for their invaluable comments on the manuscript; to Ruba Dawud, Shifaa Ghazalin, Yuval Hadar and Tehila Leitner for fly husbandry; to Ruba Dawud for her assistance with the statistical analysis; and to Christos Delidakis, Robert Duronio, Bruce Edgar, Doron Ginsberg, Gerardo Jiménez, Christian Lehner, Marco Milán, Stefan Thor, Hongyan Wang, the Developmental Studies Hybridoma Bank and the Bloomington Stock Centre for DNA constructs, antibodies, reagents and fly stocks. We thank Yael Feinstein-Rotkopf from our Faculty’s Core Research Facility for her technical assistance with microscopy. Work was supported by grants from the Israel Science Foundation (1441/20) and the Król Charitable Foundation to ZP, who is an incumbent of the Lady Davis Professorship in Experimental Medicine and Cancer Research. SBC was supported by a Bester PhD Scholarship.

## Author Contributions

SBC and ZP conceived and designed the study. SBC performed the experiments. SBC and ZP analyzed the data and wrote the manuscript.

